# A novel method to identify cell-type specific regulatory variants and their role in cancer risk

**DOI:** 10.1101/2021.11.11.468278

**Authors:** Cynthia A. Kalita, Alexander Gusev

## Abstract

**Background:** Expression quantitative trait loci (eQTLs) have been crucial in providing an understanding of how genetic variants influence gene expression. However, eQTLs are known to exert cell type specific effects, and existing methods to identify cell type specific QTLs in bulk data require large sample sizes.

**Results:** Here, we propose DeCAF (DEconvoluted cell type Allele specific Function), a new method to identify cell-fraction (cf) QTLs in tumors by leveraging both allelic and total expression information. Applying DeCAF to RNA-seq data from TCGA, we identified 3,664 genes with cfQTLs (at 10% FDR) in 14 cell types, a 5.63x increase in discovery over conventional interaction-eQTL mapping. cfQTLs replicated in external cell type specific eQTL data and were more enriched for cancer risk than conventional eQTLs. The intersection of tumorspecific QTL effects (tsQTLs) with GWAS loci identified rs4765621 and *SCARB1*, which has been previously linked to renal cell carcinoma (RCC) progression and experimentally validated in tumors.

**Conclusions:** Our new method, DeCAF, empowers the discovery of biologically meaningful cfQTLs from bulk RNA-seq data in moderately sized studies. Our study contributes to a better understanding of germline mechanisms underlying the anticancer immune response as well as cfQTLs contributing to cancer risk.

## Introduction

Genome-wide association studies (GWAS) have been instrumental in identifying a large number of genetic variants associated with risk for many diseases including cancer Scelo et al. (2017); Melin et al. (2017); Huyghe et al. (2019); McKay et al. (2017); Schumacher et al. (2018); Phelan et al. (2017); Michailidou et al. (2017). However, as the majority of GWAS associations are non-coding variants without clear function, the mechanism of action is typically unknown. Expression quantitative trait loci (eQTLs) have been instrumental in linking genetic variation to effects in gene expression Albert and Kruglyak (2015); Aguet et al. (2017); Gibbs et al. (2010); Melzer et al. (2008), recently identifying putative susceptibility genes for ovarian cancer Gusev et al. (2019a), prostate cancer Mancuso et al. (2018), and breast cancer Wu et al. (2018). Recently, it has been observed that some eQTL effects can be observed only in specific contexts and often vary across tissue and cell types Ardlie et al. (2015); Gamazon et al. (2018); Zhang et al. (2018); Schmiedel et al. (2018). In the context of cancer, the cell types in the tumor/microenvironment can have have substantially different functions Chen and Mellman (2017); Finotello and Trajanoski (2018): with CD4 and CD8 T cells driving cytotoxic anti-cancer immunity Chen and Mellman (2013; 2017), while regulatory T cells are associated with immune suppression and homeostasis Savage et al. (2013); Finotello and Trajanoski (2017). Identifying and quantifying cell type specific eQTLs in tumors is thus a critical step to understanding germline cancer mechanisms and germline-somatic interactions.

To date, cell type specific studies have been limited in size due to cost and labor associated with selecting a pure subset (ie. cell sorting) and therefore have weak power to identify QTLs Chen et al. (2016). While emerging single-cell technologies have the potential to precisely measure expression in specific cell populations, this approach remains too expensive to measure across hundreds of individuals and exhibits very sparse expression, with most genes unexpressed in a given cell. This has led to the development of bulk RNA-seq deconvolution methods which generally work by defining gene sets in reference data from a pure population and ranking the expression of these in the target sample Becht et al. (2016); Aran et al. (2017), or modelling the target sample as a mixture of signatures Gentles et al. (2015); Finotello et al. (2019); Racle et al. (2017); Li et al. (2016). These cell fraction estimates can additionally be incorporated into eQTL analyses to identify cell-fraction specific effects (cfQTLs) Geeleher et al. (2018); Wang et al. (2018a); Westra et al. (2015); Zhernakova et al. (2017); Kim-Hellmuth et al. (2020). However, by testing for an interaction effect on a very noisy outcome, such studies typically require sample sizes in the thousands to achieve adequate power (particularly for cell types present at low frequency) Westra et al. (2015).

In addition to conventional eQTLs, genetic effects on expression can be measured by quantifying the ratio of RNA-seq reads at heterozygous variants in exons. A significant departure from a 50% allelic ratio is indicative of a cis-genetic effect on expression, and referred to as allelic imbalance (AI). This approach benefits from being able to control for trans-variation, and, because it measures allelic effects within individuals and not between individuals, can harness power from read depth Cowper-Sal lari et al. (2012); Hasin-Brumshtein et al. (2014); Kasowski et al. (2010); Kukurba et al. (2014); Battenhouse et al. (2010); McVicker et al. (2013); Pastinen (2010); Timothy et al. (2012); Skelly et al. (2011). AI has also been leveraged to identify genes undergoing gene-environment interactions(Knowles et al. 2017; Moyerbrailean et al. 2016), tumor/normal regulatory differencesGusev et al. (2019b) and, recently, cell type specificity Fan et al. (2021). However, the integration of total expression and AI to detect cfQTLs has been largely unexplored.

Here we propose DeCAF (DEconvoluted cell type Allele specific Function), a method that increases power to identify cfQTLs in bulk data by combining AI and total expression signals. We use DeCAF to identify thousands of allelic cfQTLs in multiple cell types as well as tumor cells in renal cell carcinoma, which we replicate using external cell/tissue type data and link to GWAS risk variants. DeCAF identified many cfQTLs that were not observed in a conventional bulk eQTL analysis, suggesting that this framework may greatly enrich our understanding of cis regulatory mechanisms.

## 1 Results

### Overview of DeCAF for cfQTL discovery

We sought to identify cfQTL variants whose effect on gene expression varied with cell type fraction (Figure 1a). As in previous workNédélec et al. (2016); Mangravite et al. (2013); Maranville et al. (2011; 2013); Strober et al. (2019), such variants can be identified through a linear interaction model with deconvoluted (individual-level) cell fraction, where a significant interaction between genotype and cell fraction is indicative of an effect size that differs with differing cell-fractions. We introduce an analog of the interaction model for allelic imbalance (AI) data, which we combine with the conventional interaction (ieQTL) model. The ieQTL model is a linear regression of *y* ∼ *µ* + *βx* + *f* + *β*_*f*_ *x* * *f*, where *y* is total expression, *µ* is the expression mean, *x* is the genotype encoding the number of alternative alleles, *β* is the standard eQTL effect (expected to be zero under the null), *f* is the individual-level cell fraction, and *β*_*f*_ measures the cell fraction specificity. Our proposed AI model relates the number of reference and alternative allelic reads in heterozygous individuals to their cell fraction as a regression of *REF, ALT* ∼ *µ* + *α*_*f*_ *f*, where *µ* is the mean allelic fraction across individuals and *α*_*f*_ is the modifying effect of the cell-fraction *f* and our primary effect of interest. Under the null *µ* is 0.5 and deviations from 0.5 indicate an AI effect in the bulk population; likewise, a significantly non-zero *α*_*f*_ indicates the cell fraction modifies the allelic fraction effect and is indicative of a cfQTL. The parameters of the AI model can be estimated by binomial/logistic regression, but we additionally implement beta-binomial regression to accommodate read overdispersion that is common in RNA-seq data Van De Geijn et al. (2015) (see Methods). Finally, the *β*_*f*_ and *α*_*f*_ estimates (which are both oriented towards the alternative allele) are combined by meta-analysis to form the complete DeCAF test statistic. In practice, we additionally include a tumor purity term in both models to jointly estimate tumor-specific effects (tsQTLs) Geeleher et al. (2018) and determine whether a variant is acting in the tumor or the microenvironment. We note that, in contrast to some analyses of AI in individual samples, our model always tests for consistent effects in the population at a given variant, polarized to the variant allele. We tested causal variants outside the gene by using haplotype phasing (heterozygous variants were assigned the reads on their corresponding haplotype as in Gusev et al. (2019b)).

**Figure 1:**
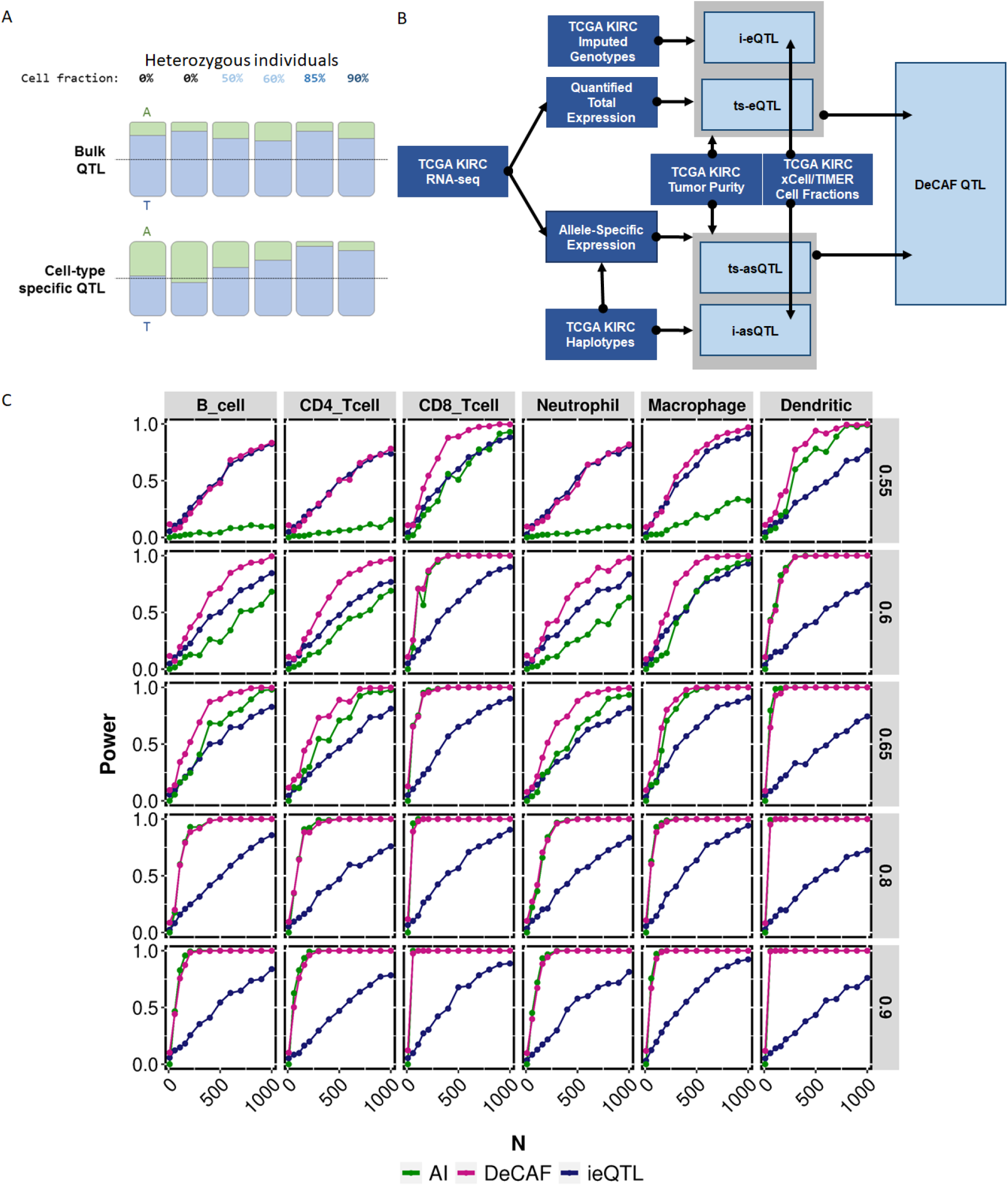
Identifying cfQTLs. A) Depiction of AI changing across increasing cell fraction. B) Flowchart showing the data entering into identifying DeCAF QTLs. Dark blue boxes represent outside data and light blue boxes represent internal calculated data. C) Comparing cfQTL testing methods for simulated data. Plot depicting the power (y-axis) for varying number of individuals (x-axis) and either ieQTL (green), 6iAI (pink), or DeCAF combined method (blue). Each column represents a cell type and each row is a different allelic fraction (effect size of AI).

Somatic copy number alterations can lead to extra variation and non-genetically driven AI. Indeed, the earliest applications of allelic imbalance in cancer were to identify copy number changes from sequenced DNA. As CNVs have been estimated in TCGA data from tumor/normal genotyping, the local CNV estimate from each individual was included in the model as a fixed effect covariate, to account for an offset in the expression due to carrying a CNV. Accounting for CNV as a covariate was well calibrated and led to only a 1.12x reduction in power (Supplement Figure S2). In the real data we additionally conservatively masked out any individuals with deep CNVs (*>*0.1 segment mean, or Log2 ratios of the tumour copy number to the normal copy number) for a given test.

### Evaluation of DeCAF in simulation

We performed wide-ranging simulations reflecting conditions found in real data and evaluated the power of the interaction eQTL, interaction AI, and DeCAF tests (see Methods). These included allelic effect size, minor allele fraction, sequencing depth, CNV, number of individuals, and read overdispersion. Under the null hypothesis where there is no effect, all of the methods were well calibrated and produced null results (Supplement Figure S1). Under the alternative hypothesis, DeCAF consistently met or outperformed the power of the conventional interaction eQTL test, requiring 0.56x fewer samples on average to reach *>*75% power (Figure 1c). In 42% of the simulated models, DeCAF achieved *>*75% power at *<*600 individuals (the typical TCGA study size) while the eQTL test did not (Figure 1c).

### Identifying a multitude of QTLs

We applied DeCAF to genotype and RNA-seq data from 503 RCC tumors in TCGA (Figure 1b) using cell fractions from xCell Aran et al. (2017) and TIMER Li et al. (2016) as well as tumor purity (previously estimated by Aran et al. (2015)) to account for differences in purity and identify tsQTLs Geeleher et al. (2018) (Figure 1b; See Methods). Across all tested cell types, DeCAF identified 5.63x more genes with significant cfQTLs than a conventional interaction QTL analysis. By cell fraction type, DeCAF identified 1753 (xCell), 1220 (TIMER) cfQTL genes and 691 (purity) tsQTL genes at 10% FDR (defined as unique genes containing a significant corresponding QTL). For comparison, conventional interaction eQTL mapping identified 310 (xCell), 218 (TIMER), and 186 (purity) tsQTL genes (Figure 2). Applying DeCAF without a cell fraction term identified a total of 3654 marginal eQTL genes, a 1.7x increase over the 2150 eQTL genes identified with the conventional eQTL linear model, suggesting that incorporating AI particularly increases power for interaction analyses where the effective sample sizes are lower. Across the cfQTL genes, we observed 42 genes with significant effects in all cell types while the rest of genes were largely observed in a single cell type (270 genes, Figure 2b) and were not substantially shared across the xCell and TIMER results. We thus focus on cfQTLs from both methods moving forward.

**Figure 2:**
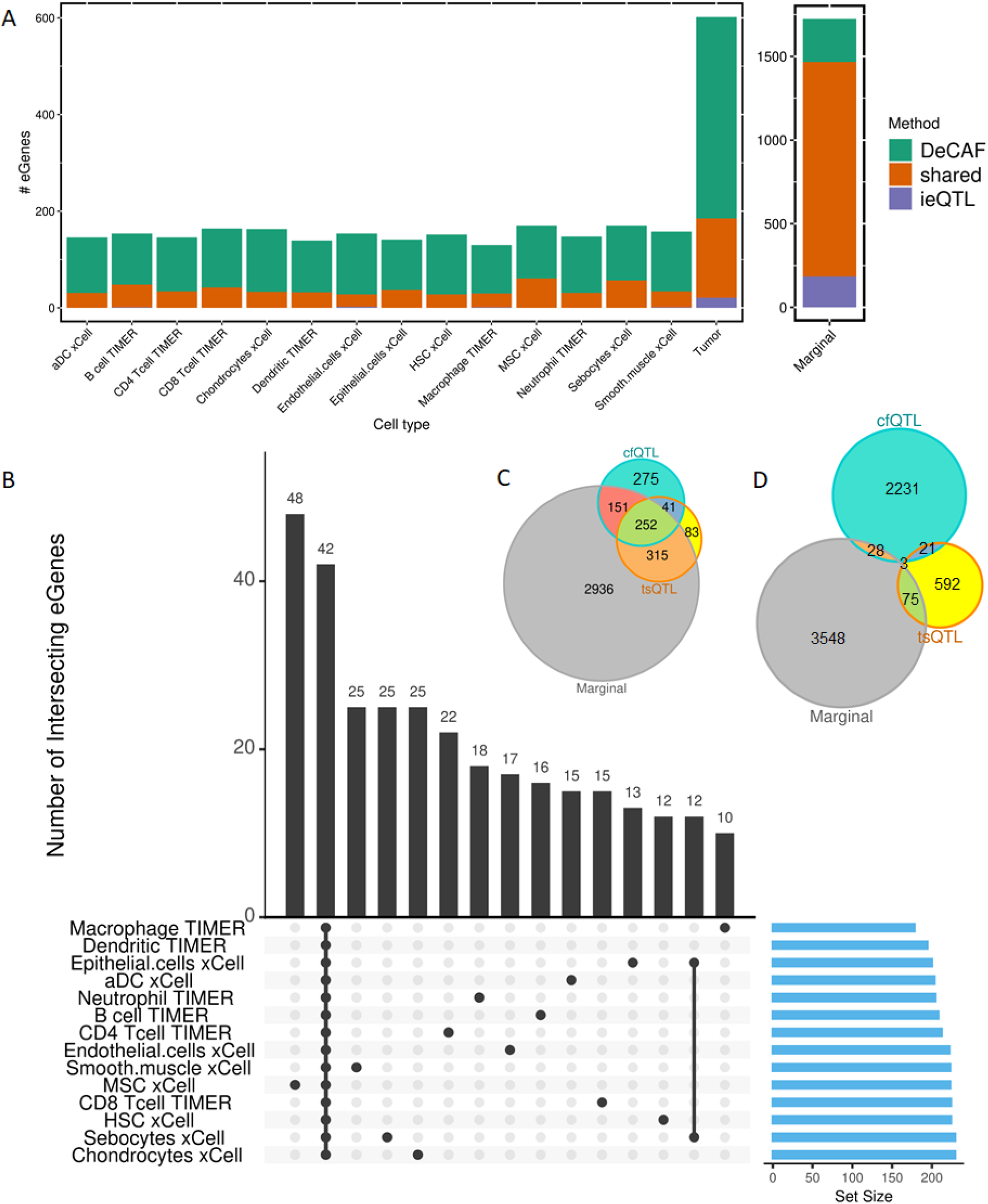
cfQTL testing methods. A) Comparing eGenes from DeCAF vs standard interaction QTL (ieQTL) methods. Plot depicting the number of significant cfQTL genes (y-axis) for each cell type (x-axis) for DeCAF only (teal), standard interaction eQTL (ieQTL) only (purple), and shared (orange) tests. B) Intersection of s8ignificant eGenes from cancer and cell fraction tests for xCell and TIMER deconvolutions. C) Overlap of eGenes from cfQTLs, tsQTLs, and marginal eQTLs. D) Overlap of significant snp:gene pairs from cfQTLs, tsQTLs, and marginal eQTLs.

Of the significant DeCAF cfQTL genes, 56% were also significant DeCAF marginal eQTL genes and 41% were also significant DeCAF tsQTL genes. In contrast, of the significant DeCAF tsQTL genes, 82% were also DeCAF marginal eQTL genes, suggesting that tumor-specific effects often manifest as apparent bulk eQTLs (although they make up a minority of all marginal eQTLs, as previously observed Geeleher et al. (2018)). Remarkably, while we saw a large portion of shared eGenes between cfQTLs, tsQTLs, and marginal eQTLs, the SNPs associated with each eGene were mostly not shared (Figure 2d). Of the significant DeCAF cfQTLs, 1% were also significant DeCAF marginal eQTLs and 1% were also significant DeCAF tsQTLs. Most strikingly, only 3 SNP:eGene pairs are shared between cfQTLs, tsQTLs, and marginal eQTLs. While differences in the lead eQTL SNP may be due to statistical noise, we further observed that for genes with an eQTL and cfQTL, 92% of SNP pairs had *r*^2^ < 0.5, suggesting that DeCAF identified largely independent genetic signals that were not detectable from conventional bulk analyses.

To provide visual intuition for the DeCAF approach, we identified genes with a significant (Spearman) correlation between cell fraction and individual allelic fraction of their corresponding cfQTL (Figure 3c). We highlight the cfQTL effect of rs1176895555 on the expression of GOLGABL1 in the context of neutrophils (*ρ* = 0.92, *p*-value = 0.001). A striking linear relationship can be observed between the neutrophil fraction in each heterozygous sample and the allele fraction, shifting from imbalance for the reference allele to imbalance for the alternative allele (i.e. a complete reversal of effect-size direction). We note that this correlation-based approach was statistically much weaker than DeCAF because it did not incorporate read count or read overdispersion, and so is used here only to provide intuition.

**Figure 3:**
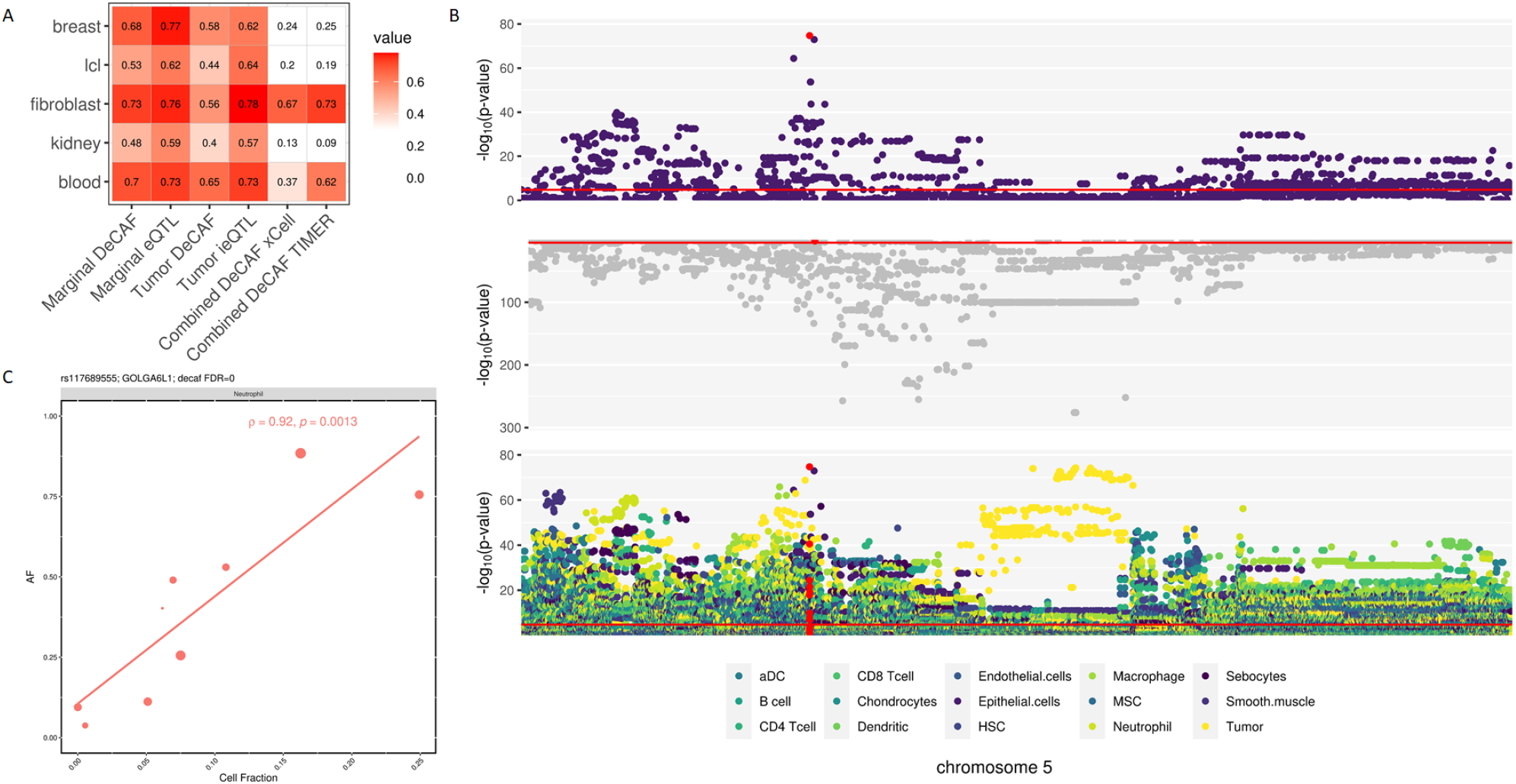
cfQTL validation. A) *π*_1_ replication of marginal, cancer, and TIMER and xCell cfQTLs in GTEx. B) ERAP2 manhattan plot with top: Epithelial cell cfQTL (most significant cell type), middle: Marginal, and bottom: all cfQTL and tsQTL significance levels. Red highlighted SNP is rs26481. C) Correlation between cell fraction (x-axis) and allelic fraction (y-axis). Point size indicates number of reads.

### Replication of DeCAF cfQTLs in external studies

Reasoning that the DeCAF cfQTLs would capture components of the tumor immune microenvironment, we sought to replicate them in external non-cancer data. First, we turned to eQTLs in the GTEx (v8 Aguet et al. (2020)), detected in bulk RNA-seq from lymphoblastoid cells (LCLs), fibroblasts, kidney, and breast tissues - selected as a sampling of potential cells/tissues in the RCC microenvironment. Across all DeCAF cfQTLs combined, we observed substantial replication across multiple GTEx tissues, with the highest replication rates in fibroblasts: *π*_1_ = 0.67 and *π*_1_ = 0.73 for xCell and TIMER-based results respectively (see Methods, Figure 4). These replication rates were comparable or higher than the replication of marginal eQTLs into GTEx data, where *π*_1_ ranged from 0.59 (kidney) to 0.77 (breast), demonstrating that cfQTLs have similar true positive rates when evaluated in the appropriate target tissue. Moreover, GTEx breast, kidney, and LCLs exhibited the lowest cfQTL replication rates (all *π*_1_ ≤ 0.25), demonstrating that cfQTLs were also specific to a target tissue (in this case fibroblast and blood), in contrast to marginal eQTLs that generally replicated in all target tissues (all *pi*_1_ *>* 0.59). Taken together, these results suggest that cfQTLs may be capturing genetic effects enriched in cancer-associated fibroblasts and depleted in organ tissues or B-lymphocytes. Within the individual cell types tested by DeCAF, we found a wide range of replication rates in GTEx tissues with no clear cell-tissue relationships, likely due to the relatively low number of cfQTLs within a given cell type or heterogeneity in the target GTEx samples (Supplement Figure S5). We note that the GTEx eQTLs did not use AI signals, further underscoring the DeCAF replications given the different statistical models employed.

**Figure 4:**
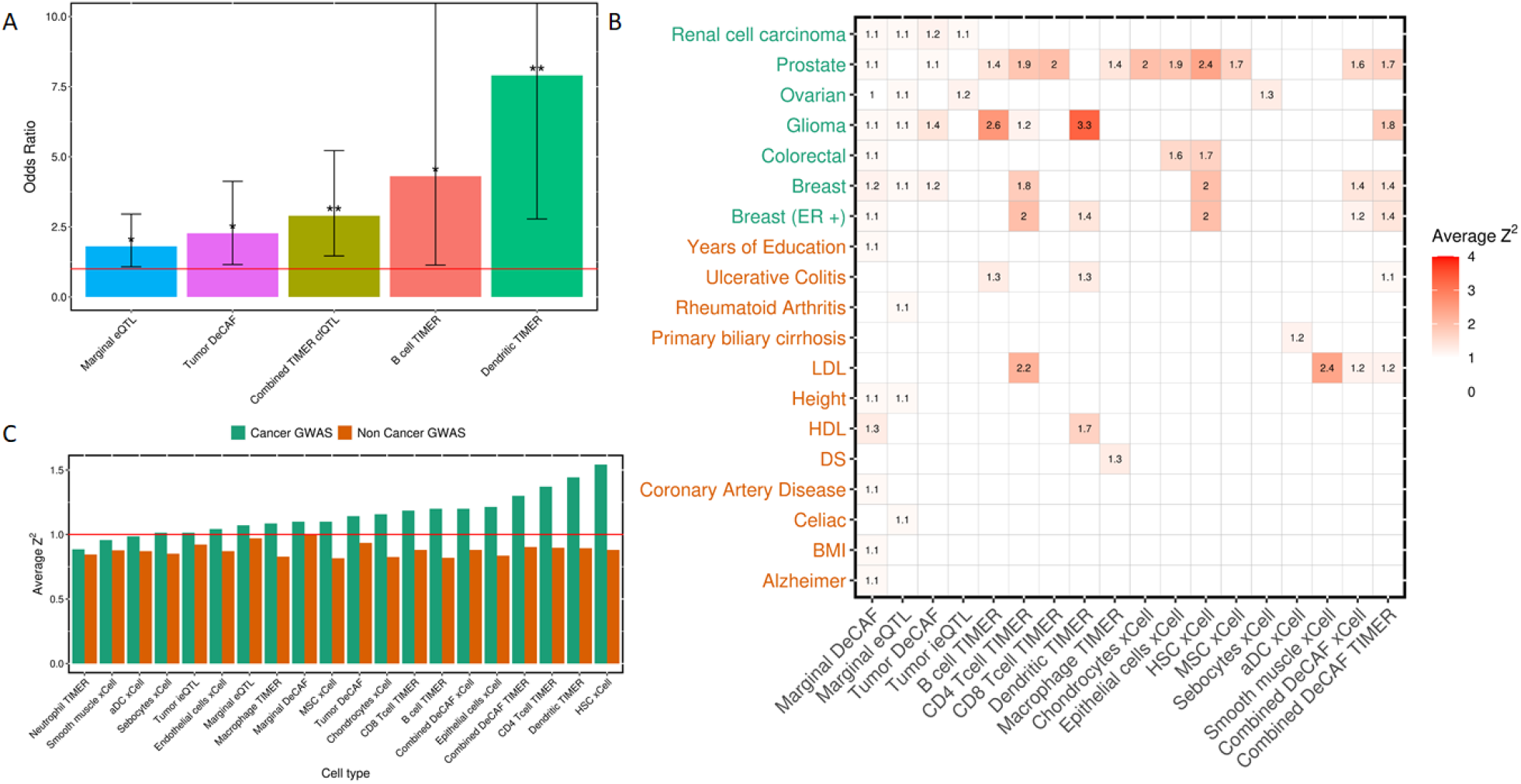
Enrichment of cfQTLs in GWAS. A) Significant enrichment of cfQTLs (FDR 20%) in RCC GWAS (*p*-value<0.001) from a fisher’s test. Two stars above the bar represents significant after Bonferroni correction, one star represents nominal significance (*p*-value*>*0.05). B) Heatmap showing the average *Z*^2^ enrichment of cfQTLs in GWAS. Insignificant results (no reported value) are based on *p*-value*>*0.05. C) Comparing average *Z*^2^ (y-axis) enrichment from cancer (green) vs non-cancer (orange) GWAS traits for each cell type (x-axis).

We carried out a second replication using cell type specific eQTL data from the BLUEPRINT consortium Chen et al. (2016). Notably, BLUEPRINT data (but not eQTLs) were used to train the xCell deconvolution scores and thus provide an apples-to-apples comparison of pure versus deconvoluted eQTLs in independent data. Across all DeCAF cfQTLs combined, replication rates were notably lower than in GTEx, with *π*_1_ in the 0.30-0.40 range (Supplementary Figure S4). As in GTEx, we again did not observe clear trends between the DeCAF cfQTL cell type and the target eQTL cell type (for example, the strongest replication was from Mesenchymal stem cells (MSCs) to CD4 T cells, with *π*_1_ = 0.60). The lower replication of BLUEPRINT versus GTEx may be explained by the relatively small sample size of the BLUEPRINT, and thus lower power to detect weak, cell type specific effects. Overall, these replications were generally comparable to the replication *π*_1_ range of 0.3-0.67 observed in a recent GTEx ieQTL analysis of similarly sized target studies Kim-Hellmuth et al. (2020), suggesting that replicating cfQTLs may be generally more challenging than marginal eQTLs.

Turning to tsQTLs, we were surprised to find substantial replication into both external eQTL studies (Figure 3). The strongest replication in GTEx was in blood (*π*_1_ = 0.65) and the strongest replication in BLUEPRINT was in neutrophils and monocytes (*π*_1_ = 0.70). Strikingly, DeCAF tsQTL replication was higher than DeCAF cfQTL in GTEx breast, LCL, kidney, and blood as well as all BLUEPRINT tissues, and was comparable to that of marginal DeCAF QTLs (Supplement Figure S4). Given the highest replication was observed in the BLUEPRINT immune cell types, we hypothesize that tsQTLs may be capturing genetic effects from or due to tumor infiltrating lymphocytes (TILs), or a mix of TILs and the tumor cell of origin. These enrichment patterns were broadly similar when testing the smaller number of conventional tumor interaction eQTLs, underscoring the robustness of this effect to detection methodology.

### Enrichment of DeCAF cfQTLs in GWAS

We next sought to quantify the relationship between cfQTLs and genome-wide association study (GWAS) variants, which may shed light on the downstream phenotypic mechanisms of cfQTLs. We analyzed the GWAS effect-size (squared Z-score) enrichments across 7 common cancer GWAS Loh et al. (2018); Melin et al. (2017); Scelo et al. (2017); Huyghe et al. (2019); McKay et al. (2017); Phelan et al. (2017); Michailidou et al. (2017); Schumacher et al. (2018). The enrichments compared significant cfQTLs to random non-significant cfQTLs as a background to account for the frequency and LD of tested variants, with significance quantified through background sampling (see Methods). Every cancer GWAS had at least one significant cfQTL enrichment (Figure 4b) and most cfQTL cell types exhibited enrichment in cancer GWAS (mean 1.16x s.e. 0.04). Notably, cfQTLs consistently showed higher enrichments than marginal DeCAF eQTLs (mean 1.10x s.e. 0.02). Compared to cfQTLs, tsQTLs generally had weaker enrichments, but were still significant in 4/7 cancers. For example, all eight cell types with significant enrichment in prostate cancer GWAS (mean 1.84x s.e. 0.12) had a higher magnitude of enrichment than the corresponding tsQTLs (mean 1.10x). As a comparison set of phenotypes, we expanded our analyses to 12 non-cancer GWAS from recent large-scale biobank studies Loh et al. (2018); Gazal et al. (2017). We observed broadly lower enrichments in these non-cancer GWAS (mean cfQTL enrichment 0.86x s.e. 0.01), with only 5/12 non-cancer GWAS traits having any significant (*p*-value<0.05) cfQTL enrichments (compared to 7/7 cancer traits), covering a broad range of trait types (Figure 4b). Overall, cancer GWAS traits had a stronger mean enrichment for every cell type than non-cancer GWAS (Figure 4c). In sum, cfQTLs are more directly relevant to cancer GWAS mechanisms than eQTLs across a wide range of cancers.

We next focused on enrichments specific to the RCC GWAS Scelo et al. (2017) where we expected more biologically plausible cell-type specific cfQTL enrichments. In the previous analysis comparing average effect sizes to non-imbalanced cfQTLs, only tsQTLs were significantly enriched for the RCC GWAS. We thus turned to a more sensitive analysis based on the enrichment of significant versus non-significant effects evaluated by Fisher’s Exact Test (see Methods). After testing multiple DeCAF FDR and GWAS *p*-value thresholds we found that a more relaxed threshold (see Methods) produced the most confident results while identifying the most commonly seen significant cell types (Supplement Figure S6a). We identified significant (Bonferoni *p*-value<0.05) enrichment for cfQTLs in (TIMER) dendritic cells (OR=7.90, *p*-value=1.86^*−*4^) and all combined TIMER cfQTLs (OR=2.89, *p*-value=1.58^*−*3^) (Figure 5a, Supplement Figure S6b), with nominal significance for cfQTLs in (TIMER) B cells (OR=4.30,*p*-value=0.02), tsQTLs (OR=2.27, *p*-value=0.01), and marginal eQTLs (OR=1.80, *p*-value=0.02). Again, marginal eQTLs were less enriched than other significant cfQTLs. Interestingly, the RCC GWAS enrichment was generally stronger for cfQTLs in individual cell types than across all cell types, suggesting that RCC GWAS variants may be enriched for specific cell types rather than generic cell type specific variants (although the statistical power was too low to identify significant differences between cell types in this analysis).

**Figure 5:**
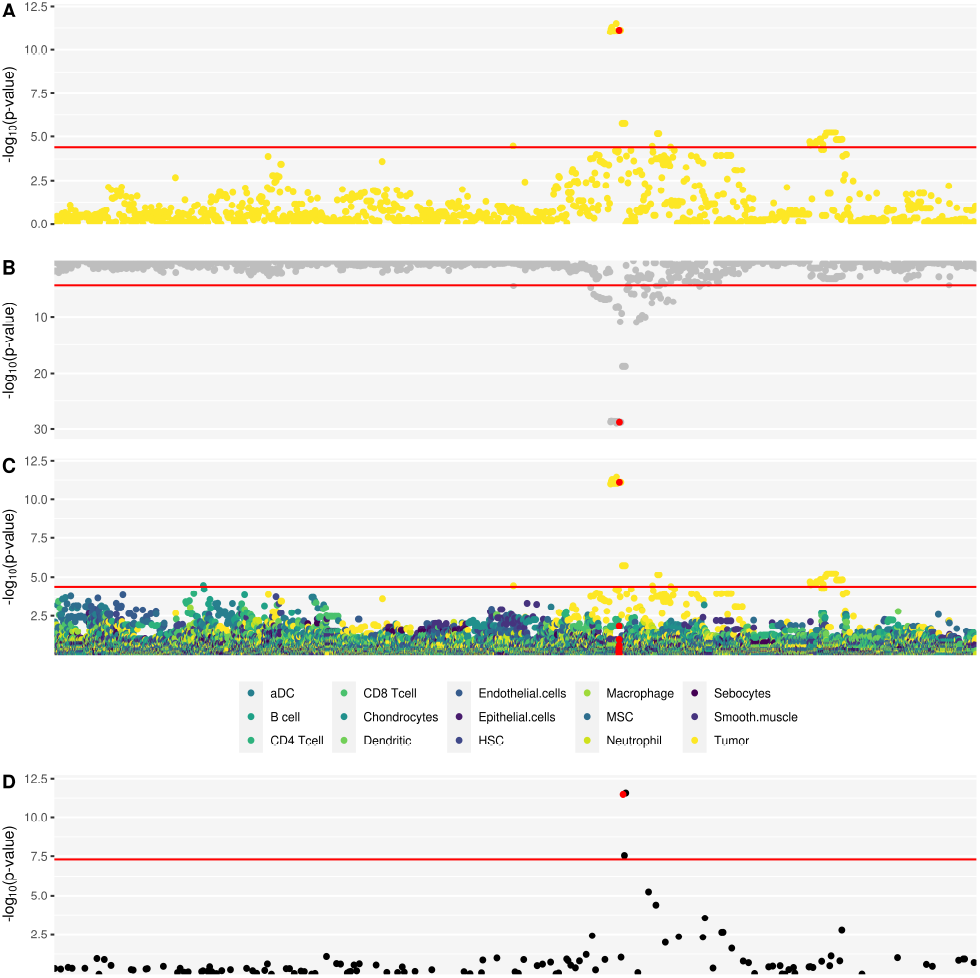
Enrichment of tsQTLs in GWAS. SCARB1 manhattan plot A) tsQTL B) Marginal C) cfQTL and D) RCC GWAS significance levels. Red highlighted SNP is rs4765621.

### Biological interpretation of cfQTL examples

Lastly, we highlight two specific examples of cfQTLs and tsQTLs that showcase the utility of this framework.

We identified a highly significant cfQTL at rs26481 and *ERAP2* in epithelial cells (FDR = 5.5^*−*69^, Figure 3b). Intriguingly, rs26481 was clearly an independent signal from the top marginal eQTL for *ERAP2* (rs2927610; FDR = 0), with an *r*^2^ of 0.0008 between the two variants. The marginal eQTL signal also coincided with a highly significant tsQTL, indicative of two independent genetic mechanisms (microenvironment and tumor) operating through *ERAP2*. Interestingly, a previous study found that ERAP2 expression was limited to epithelial cells in normal tissue, and loss of this enzyme was found in 86% of tumors Fruci et al. (2008). This loss of *ERAP2* was also significantly linked to lack of HLA class I molecules Fruci et al. (2008). In renal cancers, *ERAP2* was found to be frequently downregulated compared to normal tissue, and potentially involved in immune escape mechanisms Stoehr et al. (2013). The low expression of *ERAP2* has also been associated with poor cancer prognosis Compagnone et al. (2019).

Intersecting the DeCAF results with kidney cancer GWAS data revealed a tsQTL at rs4765621 (FDR = 6.02^*−*8^) that colocalized with a genome-wide significant significant risk variant (*p*-value = 3.33^*−*12^) for *SCARB1* (Figure 5a). The same variant was also a marginal eQTL for *SCARB1* but is not an eQTL in the normal Gusev et al. (2019b), strongly supporting its tumor-specific effect (Figure 5). Previous studies have found that inhibiting *SCARB1* (a HDL cholesterol receptor) could kill and stop proliferation of clear cell renal cell carcinoma (ccRCC) cells Riscal et al. (2021). A previous study Gusev et al. (2019b) thoroughly validated the effect of rs4765623 on *SCARB1*, a SNP in perfect LD with the variant identified as a tsQTL in our study. rs4765623/*SCARB1* exhibited tumor-specific enhancer activity and was validated in a 786-O cell line, with the C risk allele enriched for chromatin activity, consistent with it’s expression increasing effect in our study.

## Discussion

We presented DeCAF, a novel method for identifying cell type specific QTLs by harnessing signals from both total and allelic expression. No methods currently exist that leverage AI for cell type specific discovery, relying solely on QTL signals. By drawing additional power from AI (both due to it’s count-based nature, and it’s internal control for artifacts), DeCAF enables well-powered cfQTL analyses of medium-size QTL studies that would not have sufficient power for conventional tests. DeCAF directly models multiple potential cancer-specific confounders including tumor purity and CNVs, and can distinguish whether a QTL is functional in the tumor, microenvironment cell type, or both.

We applied DeCAF to TCGA data to perform the first cfQTL mapping effort in tumors. DeCAF identified 3664 significant cfQTL genes across all cell types, 5.63x more than was found using conventional ieQTL mapping (consistent with simulations). DeCAF similarly identified 3.72x more tsQTLs than the conventional approach, thus being a powerful method for identifying tumor-specific effects even in the absence of deconvoluted data. Surprisingly, DeCAF tsQTLs had the higher replication levels in independent eQTL data. This high replication, particularly in sorted immune cells, suggests tsQTLs may capture QTL effects in the context of tumor immune infiltration. Though we caution that the tsQTL analysis was substantially better powered than any other individual cfQTL analysis due to the higher variance and abundance of pure tumors, which likely impacts the relative replication performance. As little is known about the relevance of cell types in the tumor microenvironment to cancer risk, we integrated DeCAF data with cancer GWAS variants. We observed significant and cell type specific enrichment of cfQTLs with GWAS risk variants across all cancers evaluated; which was substantially higher than in non-cancer GWAS. By looking at the overlap of GWAS variants with cfQTLs and cancer QTLs, we were able to characterize their function and connect them to target genes in a specific cellular context. In sum, we found that DeCAF can identify thousands of cfQTLs from bulk RNA-seq, these cfQTLs replicated significantly in independent eQTL data (particularly tsQTLs), and were more enriched for GWAS risk than conventional eQTLs.

Other methods to detect cell type specific eQTL effects from bulk tissue data have recently been applied. GTEx Kim-Hellmuth et al. (2020) and Westra et al. Westra et al. (2015) took the approach of identifying ieQTLs using a standard linear model with a cell fraction - genotype interaction term. The GTEx study evaluated concordance with allelic fractions in the same individuals as a replication, but did not utilize the full power of cell-type AI for discovery. As such, these approaches were generally underpowered for typical, moderately sized eQTL studies. For example, Westra et al. showed very low power below N=1000 (N<1000 represents the vast majority of QTL studies), and the GTEx analysis identified <80 ieQTL genes per cell type-tissue pair on average. Our approach also shares some similarities to recently proposed methods that leverage AI in other contexts. EAGLE (Knowles et al. 2017) is a method to identify gene-environment interactions through AI mapping, which in principle could be extended to quantitative cell-fractions as “environments”. However, EAGLE only identifies a gene level effect rather than specific QTLs, and is thus challenging to integrate with external data such as GWAS. The recent method BSCET Fan et al. (2021) leveraged AI with deconvoluted cell types, but is similarly intended for analysis of transcribed SNPs rather than cfQTLs, and does not model the count nature of allelic data nor incorporate total expression. DeCAF is thus the first method to integrate total and allelic expression together for powerful cfQTL discovery at small to moderate sample sizes.

While DeCAF has made great strides in identifying cfQTLs, it has some limitations. First, DeCAF has limited power to detect cfQTLs in rare cell types (ie. low cell fractions or cell fraction variance). Lower power for rare cell types is also a limitation of direct single-cell QTL analyses and motivates larger reference and target studies. Second, DeCAF is inherently dependent on the quality of the deconvolution and cannot test cell types that are not in the reference data. The majority of existing deconvolution methods (including TIMER and xCell) calculate cell fraction scores and not direct percentages. As a consequence, DeCAF cfQTL effect sizes should be interpreted with caution, because they will be influenced by (a) the scale of the deconvoluted score; (b) the differing uncertainty in the deconvolution of different cell types; (c) and the power to detect an effect in rare cell types. While most deconvolution methods have been shown to replicate well in pure bulk cell types Becht et al. (2016); Aran et al. (2017); Gentles et al. (2015); Finotello et al. (2019); Racle et al. (2017); Li et al. (2016), they continue to be in active development. DeCAF can be applied to any deconvolution framework or score and so will benefit from improved methodologies in the future. Using DeCAF to improve the cell type deconvolution itself (i.e. by maximizing cfQTL discovery) is thus a compelling future direction. Third, we elected to take the conservative approach and remove individuals with high CNV values from the analysis. In principle, DeCAF could be extended to model both cell type and CNV specific interactions using matched allelic data from DNA sequencing, but this remains an open problem. Fourth, AI cannot be used for trans-QTL discovery, as the tested variant must be in phase with the allelic variants, and thus DeCAF’s power advantages are limited to the study of cis-eQTLs. Lastly, although DeCAF cfQTLs broadly replicated in external data at comparable rates to prior studies, we generally did not find that a given DeCAF cell type replicated best in the matching target cell type. This lack of cell type consistent replication may be explained by many factors including: differences in cell fractions influencing DeCAF power, differences in the target study sizes influencing replication power, and potential differences between AI signals in the discovery and eQTL signals in the replication. Understanding the relationship between cfQTLs detected in heterogeneous cell populations and cell-type specific QTLs detected in pure cell types thus continues to be an open question of great interest.

Although DeCAF was applied to RNA-seq data in this work, it is directly amenable to identifying cis-regulatory effects in other molecular phenotypes and contexts where sequencing read data is available. In the case of emerging single-cell QTL studies, DeCAF can be applied to identify cell-type specific QTLs using the directly estimated cell fractions. Likewise, other measures of cell state, which may span multiple cell types, could be incorporated as interaction terms in the DeCAF framework. Beyond RNA-seq, cfQTLs can be investigated via DeCAF in other molecular measurements such as tumor ATAC-seq data from TCGA Corces et al. (2018), which would shed light on cell type and tumor specific mechanisms of DNA accessibility. DeCAF could be extended to estimate effects in multiple cell types jointly, through the inclusion of multiple interaction terms, although the potential for over-fitting and autocorrelation across cell fractions needs to be carefully considered. Beyond cell fraction and tumor purity, any continuous score can be used in DeCAF, including polygenic risk scores capturing germline risk burden Shi et al. (2020), integrated clustering of molecular cancer subtypes, and emerging immune subtype definitions Thorsson et al. (2018). DeCAF thus provides a framework for broad analyses of context-specific QTLs in tumor and non-tumor data.

## Methods

### Statistical model to detect cfQTLs

The conventional QTL-based approach defines *y* as a per-individual vector of total expression, *f* as the corresponding cell fraction estimate for a given cell type, and *x* as the genotype; and the models *y* = *µ*+*βx*+*β*_*f*_ *x* * *f* . The cfQTL effect, *β*_*f*_, is then estimated by typical linear regression Nédélec et al. (2016); Mangravite et al. (2013); Maranville et al. (2011; 2013); Strober et al. (2019). To test cell type specific AI, we first restrict to individuals that are heterozygous for a variant in the target gene for which reads have been allelically assigned (we refer to this as the “functional SNP”). For this sub-population, *f* is again defined as the vector of cell fraction estimates, but instead of *x* we introduce *π*, the population-level allelic fraction for heterozygous carriers of the functional SNP. Unlike conventional allelic imbalance tests that evaluate one individual at a time internally, we test for a consistent trend of cell fraction and imbalance across the population. We then model *π* = *π*_*f*_ *f* + *π*_0_(1 − *f*) where *π*_*f*_ is the allelic fraction in the focal cell type and *π*_0_ is the allelic fraction in the rest of the cell types. While we do not observe the *π* value, for each individual we see *REF, ALT* read counts that are sampled from their *π*. The cfQTL is then estimated by binomial regression of *REF, ALT* ∼ *µ* + *β*_*f*_ *f* where *µ* captures the mean allelic fraction in the population and *β*_*f*_ captures the additional fraction-specific effect. When *µ* is significantly different from 0.5 this variant is exhibiting population-level AI, and when *β*_*f*_ is significantly different from 0, this variant is also exhibiting cell type specific AI. Functional (read carrying) SNPs were tested directly. For distal SNPs, the read counts for each allele were computed as the sum of functional SNP reads along the respective haplotype (as shown in previous work Kumasaka et al. (2016); Van De Geijn et al. (2015)). To account for overdispersion that is common in molecular dataVan De Geijn et al. (2015), we leveraged a beta-binomial regression, with overdispersion estimated for each individual across all heterozygous reads (see below for tumors). Finally, we combined the eQTL-interaction and cfQTL tests by Stouffer’s method (these tests are independent and so can be combined); we refer to the combined estimate as the DeCAF test statistic.

### Statistical model to account for tumor variation

The basic model described above allows for estimation of cfQTLs in normal expression, and we additionally extend this model to account for potential biases due tumor heterogeneity and somatic alterations. First, tumors are a mix of normal and cancer cells which introduces additional variance into the expression and could create a false relationship with a given cell type if it is correlated with the tumor fraction. To account for this, we introduce an additional term corresponding to tumor purity into the AI model, and an interaction term corresponding to the SNP-purity interaction into the eQTL model (as previously proposed in ref.Geeleher et al. (2018)). This extension improves power for cfQTLs by accounting for tumor-specific variance. In addition, tumor-specific QTLsGeeleher et al. (2018) can be inferred by testing for a non-zero effect size on the purity term.

Second, somatic copy number alterations can lead to extra variation and non-genetically driven AI. Indeed, the earliest applications of allelic imbalance in cancer were to identify copy number alterations (CNVs) from sequenced DNA. In the AI component of the DeCAF model, we estimate the beta-binomial overdispersion parameter for each CNV region in an individual separately, to account for CNV-specific variance. We additionally investigated three approaches to account for extra variance due to a significant CNV: (i) excluding variants in a CNV in an individual from the analysis entirely; (ii) including the per-individual CNV estimate in the model as a fixed effect covariate, to account for an offset in the expression due to carrying a CNV; (iii) including a random effect term for all CNV carriers to account for extra variance in expression without a consistent direction. These models make different assumptions about the CNV architecture and we selected the best performing model empirically based on the number of cfQTLs identified and their reproducibility in external data.

### Simulation framework

We investigated multiple AI-based approaches and modelling assumptions in simulation. Parameters tested for impact on the performance of these tests included: minor allele frequency (MAF); baseline (0.5) and cell type specific effect size (*AF*); read depth (*D*, sampled from a

Poisson with fixed mean); number of individuals (*N*); and cell fraction percentages (*f*, sampled from a uniform distribution or from real data). To generate a single AI individual, a *π* is defined as 0.5(1 − *f*) + (*π*_*f*_ * *f*), ALT reads were drawn from *X* ∼ BetaBin (*π, N*) with fixed overdispersion parameter, and REF reads were computed as *D* − *ALT* . Hardy-Weinberg equations were then used to generate the expected number of heterozygous AI individuals based on the MAF. To simulate CNVs, a fraction of individuals were sampled as being carriers and additional variance terms are introduced. In the fixed-effect model, being a CNV carrier adds a constant factor to the *π* (i.e. one allele is always amplified), whereas in the random effect model, one allele is randomly amplified or deleted for computing the *π*. Finally, to simulate total expression for the conventional eQTL models, quantitative phenotypes were sampled from a linear model *y* = *σ*_*NF*_ * *NF* (*σ*_*effect*_ * *SNP* * *f*) + *σ*_*CNV*_ * *CNV* + *ε*. The overall simulation was performed 500 times and power for each test was defined as the number of simulations in which that test produces an association with p<0.05/20,000 (where 20,000 is approximately the number of genes to be tested).

Wherever possible, we identified simulation parameters from the real data: mean over-dispersion was estimated as 0.0263 from tumor RNA-seq in TCGA; QTL effect sizes are set to 0.04, matching the average eQTL variance explained in real data Ng et al. (2019); normal fraction (NF) effect sizes were set to 0.0269, matching the average variance in expression explained by NF from real data; *σ*_CNV_ was set to 0.012 to match the variance in CNVs in real data.

### Deconvoluted cell fractions in TCGA data

TCGA is a rich resource for tumor RNA-seq data which has been deconvoluted into cell types by multiple published methods: xCell Aran et al. (2017) (64 cell types), which defines gene sets in a pure population and ranks the expression of these in the sample; TIMER Li et al. (2016) (six tumor-infiltrating immune cell types including B cells, CD4 T cells, CD8 T cells, neutrophils, macrophages, and dendritic cells), which uses a signature gene matrix from bulk expression; and others Wang et al. (2018b); Roman et al. (2017); Charoentong et al. (2017). We downloaded xCell Aran et al. (2017) and TIMER Li et al. (2016) cell fraction data for individuals with KIRC data from TCGA. To further improve power for testing in the xCell results, we focus on using cell types with a cell fraction score with an interquartile range *>*0.1 and cell types expected to be found in kidney tissue (Supplement Figure S3). Previous estimates of tumor purity Aran et al. (2015) were additionally used to account for differences in purity and identify tsQTLs Geeleher et al. (2018) (Figure 1b).

### Allele-specific quantification

We applied DeCAF to genotype and RNA-seq data from 503 TCGA RCC tumors from the KIRC study (Figure 1b). Germline genotype data and RNA-seq BAMs were downloaded from the NCI Genomic Data Commons. Germline genotype data was imputed and phased through the Michigan Imputation Server. Total quantified and normalized gene expression was accessed as previously described Broad Institute TCGA Genome Data Analysis Center (2016). RNA-seq reads were integrated with the genotype data and run through the WASP allelic mapping pipeline to account for potential mapping biases. In brief, for each read containing a polymorphism a new “proxy” read was generated with the alternative allele and mapped to the genome; any reads with proxies that did not map to the same location were then excluded from the analysis. This conservative approach removes reads with potential allele-specific mapping biases. Finally, allelic counts at individual variants were quantified using GATK Van der Auwera and O’Connor (2020) ASE quantification and assigned to the phased haplotypes. To process the TCGA data efficiently, we implemented DeCAF in a self-contained and reproducible pipeline on the Cancer Genomics Cloud, creating a workflow that can then be run on any data source (see Web Resources).

### cfQTL discovery

For QTL discovery, for each expressed gene and cell type, we tested each common (MAF greater than 1%) germline SNP in the cis-locus of the gene (100kb around the TSS) using each described method. Significant cfQTLs in each cell type were determined by identifying the most significant variant for each cell type and gene, applying a genel-level Bonferroni correction, followed by a Benjamini-Hochberg False Discovery Rate correction across all genes within a cell type. The same testing and multiple test correction procedure was applied to identify significant marginal eQTLs in the bulk data.

### Testing for variant enrichment

We calculated multiple enrichment and replication estimators for cfQTLs. First, we estimated *π*_1_ replication statistics (i.e. the fraction of associations that are expected to be “non-null”) using the qvalue package Storey and Tibshirani (2003) and (FDR<5%) for the significance threshold for the DeCAF QTL input. Secondly, we conduct a Fisher’s Exact Test for enrichment as follows. A contingency table was constructed where positive-positive was defined as the number of SNPs with significance in both DeCAF (FDR<20%) and GWAS (*p*-value<0.001). Adjustment was based on the Bonferroni method. Finally, the average *Z*^2^ was calculated as follows. We calculated the average *Z*^2^ (Z score of the SNPs in the category of interest) for significant DeCAF cfQTLs overlapping the category of interest. We then calculated the average *Z*^2^ of a randomly sampled (100X) null and took the ratio of the significant average *Z*^2^ over the null as the measure of enrichment.

## Data Availability

Software implementing STRATAS DeCAF can be found at https://github.com/cakalita/stratAS/tree/DeCAF. Workflow generating the allelic counts can be found on the Cancer Genomics Cloud (CGC) Lau et al. (2017) https://cgc.sbgenomics.com/u/cakalita/quickstart/apps/#cakalita/quickstart/stratas_v3.

## Acknowledgements

A.G. and C.K. were supported by R01CA227237, R01CA244569, and the Doris Duke Charitable Foundation. C.K. was supported by the Louis B. Mayer Foundation and the Claudia Adams Barr Foundation.

The Seven Bridges Cancer Research Data Commons Cloud Resource has been funded in whole or in part with Federal funds from the National Cancer Institute, National Institutes of Health, Contract No. HHSN261201400008C and ID/IQ Agreement No. 17×146 under Contract No. HHSN261201500003I and 75N91019D00024.

## Supplement

Table S1: **DeCAF sigificant SNPs**. Listed for each significant (FDR 10%) SNP is the SNP, gene, cell-type, Z score, *p*-value, and FDR corrected *p*-value.

*See attached file*

**Figure S1:**
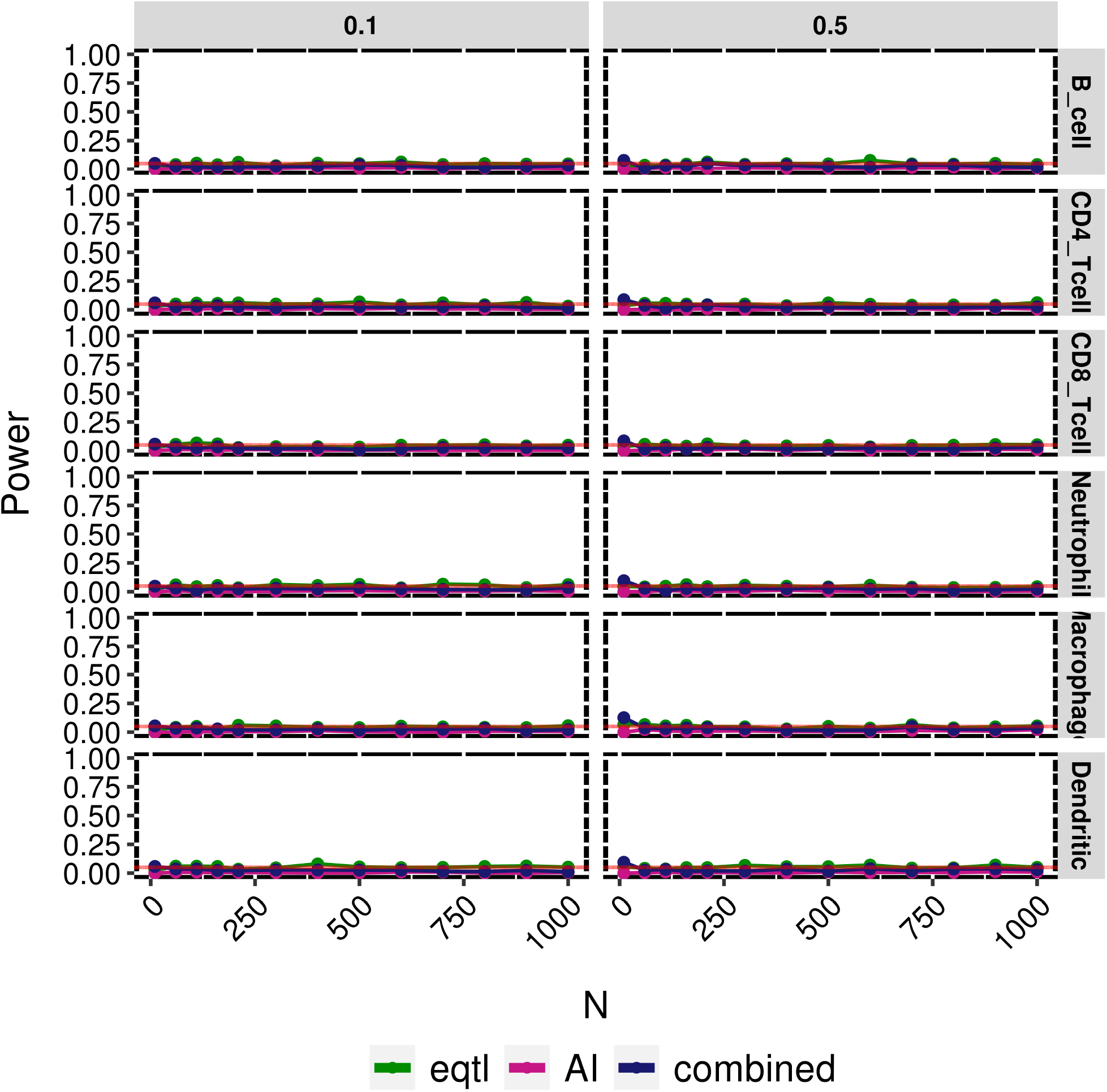
Null cfQTL models for simulated data. Plot depicting the power (y-axis) for varying number of individuals (x-axis) and either eQTL (green), AI (pink), or DeCAF combined method (blue) at a null allelic fraction (effect size of AI).

**Figure S2:**
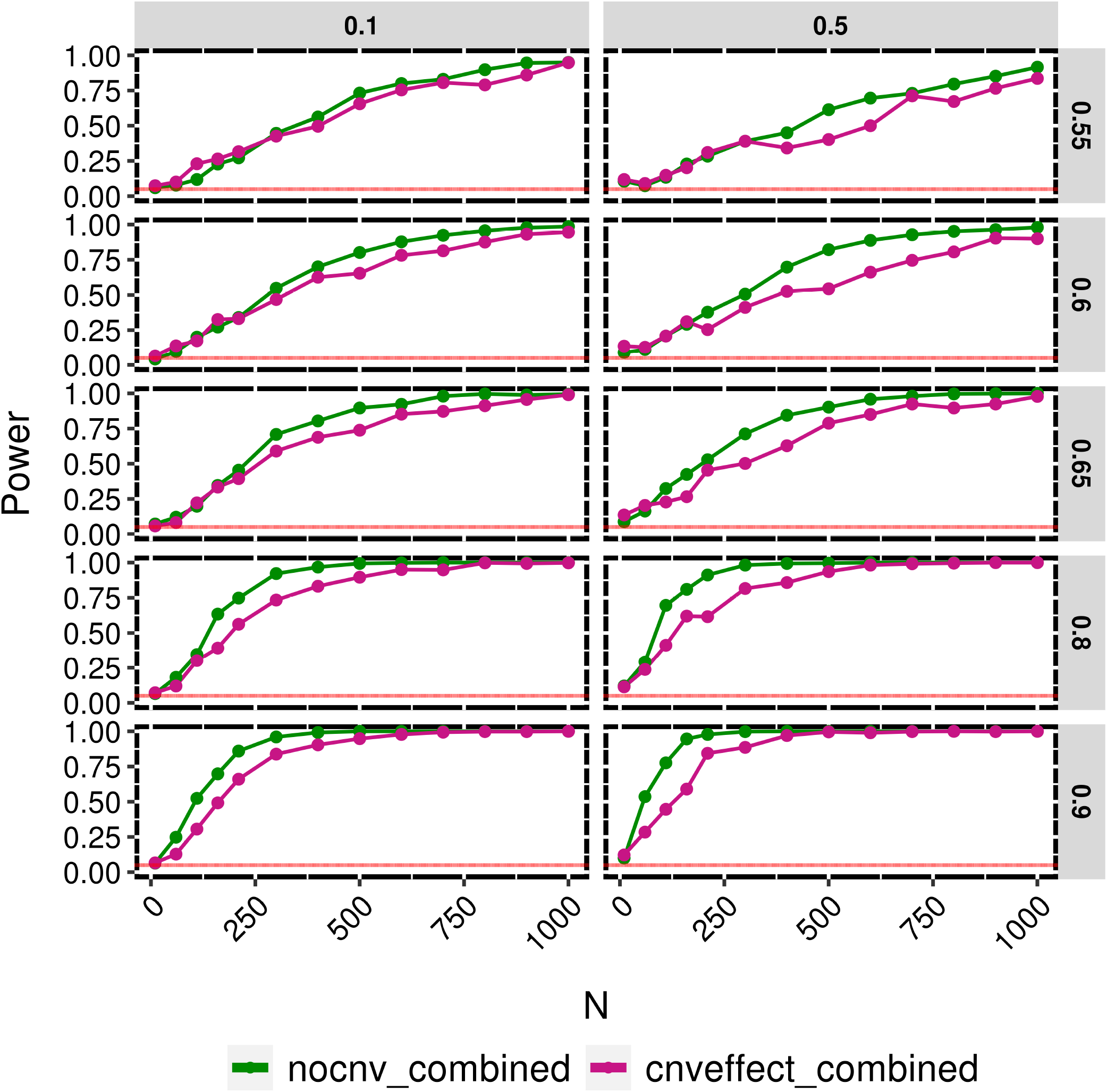
Comparing cfQTL models with or without CNV for simulated data. Plot depicting the power (y-axis) for a varying number of individuals (x-axis) and either no CNV in the DeCAF testing model (green) or CNV in the DeCAF testing model (pink). Each column represents a different MAF and each row is a different allelic fraction (effect size of AI).

**Figure S3:**
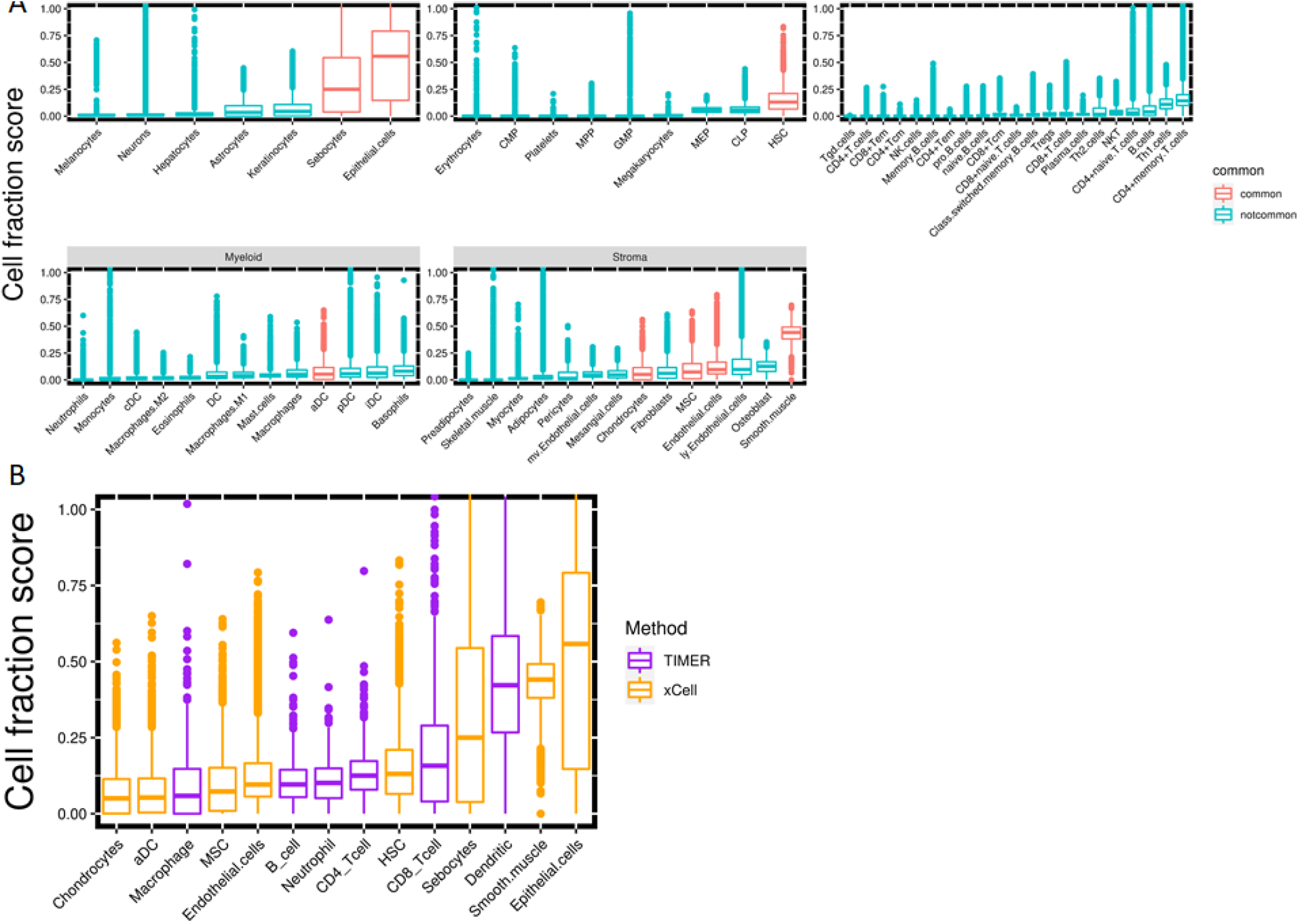
Deconvoluted cell fractions. A) Plot depicting the range of calculated cell fractions from xCell and TIMER deconvolution methods for TCGA KIRC tumors. B) All cell-type fractions are shown for xCell and a cell fraction score inter quartile range (IQR) *>* 0.1 are in pink.

**Figure S4:**
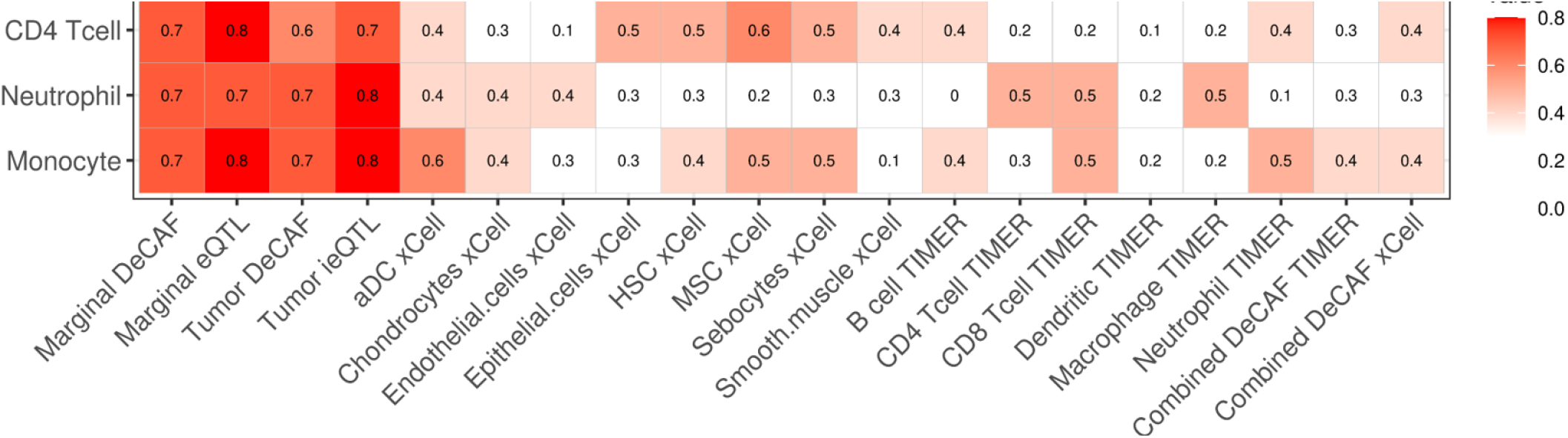
BLUEPRINT Replication. A) Average *Z*^2^ replication of marginal, cancer, and TIMER and xCell cfQTLs in BLUEPRINT. Insignificant results (crossed out values) are based on *p*-value*>*0.05. B) *π*_1_ replication of marginal, cancer, and TIMER and xCell cfQTLs in BLUEPRINT.

**Figure S5:**
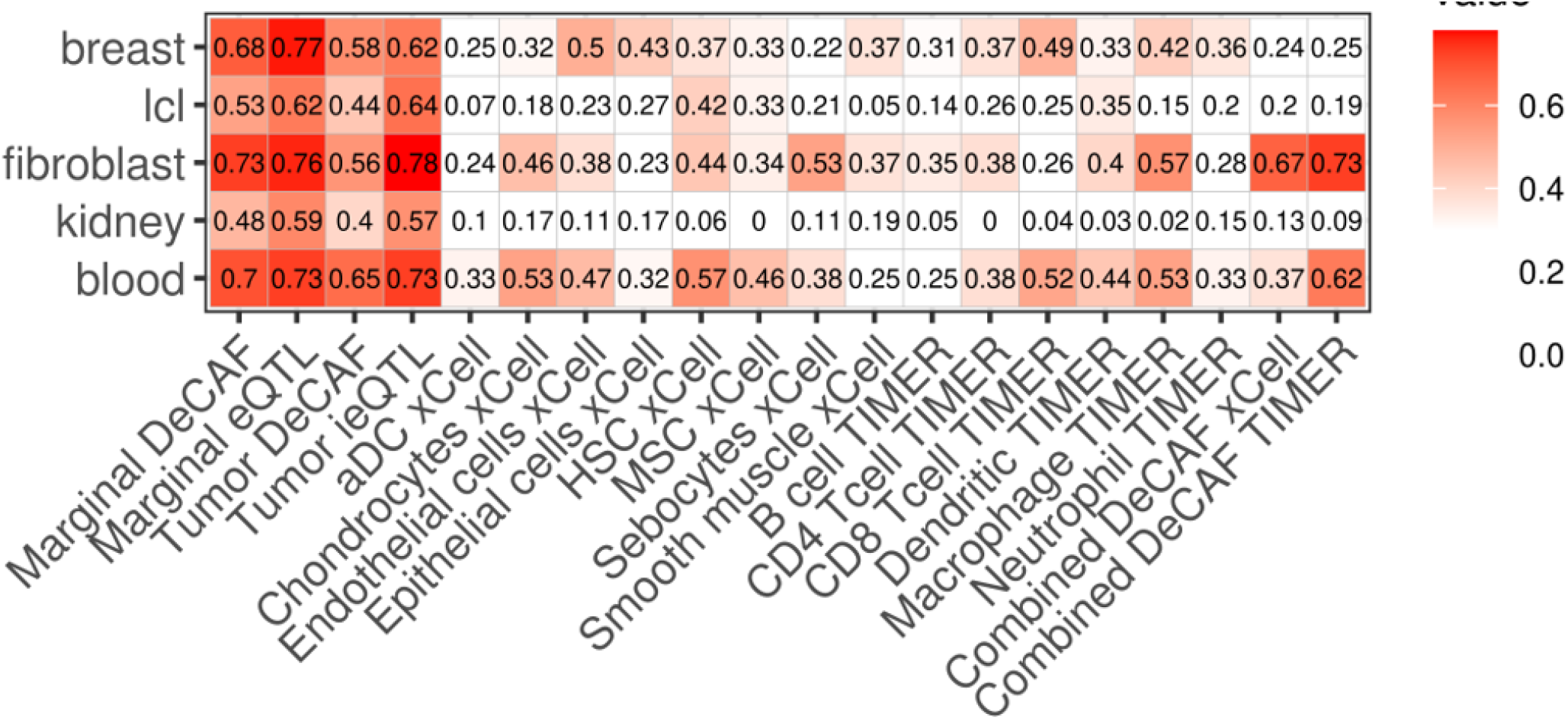
GTEx Replication. A) Average *Z*^2^ replication of marginal, cancer, and TIMER and xCell cfQTLs in GTEx. Insignificant results (crossed out values) are based on *p*-value¿0.05. B) *π*_1_ replication of marginal, cancer, and TIMER and xCell cfQTLs in GTEx.

**Figure S6:**
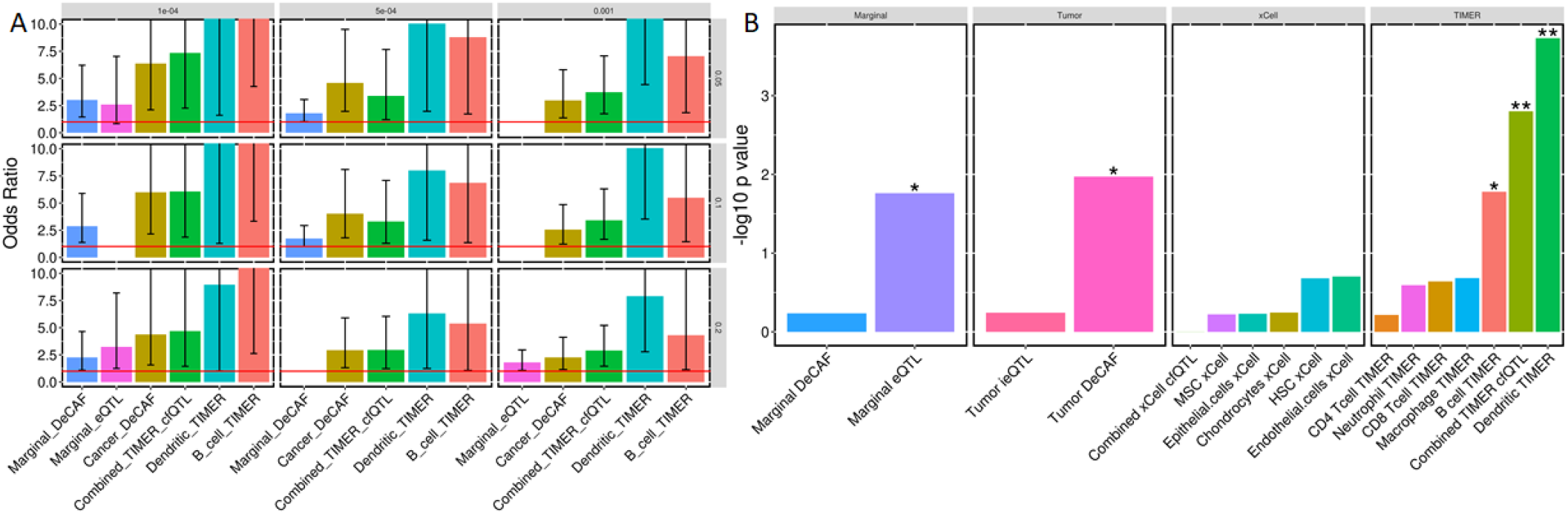
Enrichment of cfQTLs in GWAS. A) Significant enrichment of cfQTLs (FDR thresholds rows) in RCC GWAS (*p*-value thresholds columns) from a fisher’s test. B) Significant enrichment of cfQTLs (FDR 20%) in RCC GWAS (*p*-value*<*0.001). Two stars above the bar represents significant after Bonferroni correction, one star represents nominal significance (*p*-value*>*0.05).

